# Integrative Omics Reveals Subtle Molecular Perturbations Following Ischemic Conditioning in a Porcine Kidney Transplant Model

**DOI:** 10.1101/2021.04.23.441109

**Authors:** Darragh P. O’Brien, Adam M. Thorne, Honglei Huang, Elisa Pappalardo, Xuan Yao, Peter Søndergaard Thyrrestrup, Kristian Ravlo, Niels Secher, Rikke Norregaard, Rutger J. Ploeg, Bente Jespersen, Benedikt M. Kessler

**Author notes:** These authors contributed equally (Darragh P. O’Brien, Adam M. Thorne).

## Abstract

**Background:** Remote Ischemic Conditioning (RIC) has been proposed as a therapeutic intervention to circumvent the ischemia/reperfusion injury (IRI) that is inherent to organ transplantation. Using a porcine kidney transplant model, we aimed to decipher the subclinical molecular effects of a RIC regime, compared to non-RIC controls.

**Methods:** Kidney pairs (n = 8+8) were extracted from brain dead donor pigs and transplanted in juvenile recipient pigs following a period of cold ischemia. One of the two kidney recipients in each pair was subjected to RIC prior to kidney graft reperfusion, while the other served as non-RIC control. We designed a modern integrative Omics strategy combining transcriptomics, proteomics, and phosphoproteomics to deduce molecular signatures in kidney tissue that could be attributed to RIC.

**Results:** In kidney grafts taken out 10 h after transplantation we detected minimal molecular perturbations following RIC compared to non-RIC at the transcriptome level, which was mirrored at the proteome level. In particular, we noted that RIC resulted in suppression of tissue inflammatory profiles. Furthermore, an accumulation of muscle extracellular matrix assembly proteins in kidney tissues was detected at the protein level, which may be in response to muscle tissue damage and/or fibrosis. However, the majority of these protein changes did not reach significance (p<0.05).

**Conclusions:** Our data identifies subtle molecular phenotypes in porcine kidneys following RIC and this knowledge could potentially aid optimization of remote ischemic conditioning protocols in renal transplantation.

## BACKGROUND

Organ donation and transplantation inevitably involve periods of both warm and cold ischemia, due to diminished or restricted blood flow, which deprives the donor organ tissue of the oxygen required to maintain cellular metabolism. Subsequent reperfusion at time of implantation of the ischemic graft results in tissue damage including several molecular hallmarks, such as hypoxic stress and production of reactive oxygen species, a rapid reduction of ATP levels, intracellular acidosis, and diminished nutritional supplies [1]. Combined, these insults lead to the activation of apoptotic and necrotic pathways, and if uncorrected, to organ dysfunction, rejection, and eventually graft fibrosis [2]. Acute kidney injury (AKI) induced by ischemia and subsequent reperfusion is called delayed graft function in the clinical transplant setting and leads to the need of dialysis as well as a risk of poorer transplant outcome [3]. Remote ischemic conditioning (RIC) has been proposed as a therapeutic modality to mitigate the effects of ischemia/reperfusion injury (IRI) in organ transplantation [4, 5]. RIC is a simple procedure which creates short and repetitive cycles of non-damaging ischemia, usually applied to a leg or arm, which may reduce damaging effects caused by IRI in the target organ. RIC has shown various outcomes in pre-clinical studies in both kidney and other organs such as lung, liver, small intestine and in a variety of situations and species [6-12]. Several pre-clinical studies have demonstrated a reduction in IRI and improved early graft function in the recipient [13-16]. Regarding kidney transplantation and AKI, a protective effect of RIC in rodent renal models has been described, however the transition into the human setting has been challenging [15, 17]. Our previous work in a porcine model of kidney transplantation suggested a beneficial effect of RIC on early graft perfusion and function evidenced by a significantly improved renal plasma perfusion and glomerular filtration rate (GFR) [18]. However, when RIC was applied to kidney transplant patients in a clinical trial (CONTEXT), no clinical benefit was observed, although the trial did reveal distinct subclinical effects on kidney transplants at the molecular level [17, 19, 20]. We have therefore revisited our RIC study carried out in pigs that had shown potential positive effects of RIC which stimulated us to design the clinical CONTEXT study. Using samples obtained from our original porcine RIC model [18], we have now studied whether we could detect any proteomic and transcriptomic changes in recipient tissue using advanced quantitative mass spectrometry and RNA sequencing. In this study, we describe changes in molecular profiles comparable to the ones observed in human kidneys subjected to RIC.

## METHODS

### Transplant model

The effects of RIC versus non-RIC controls in porcine kidney transplants were evaluated in a paired randomized study, the design of which was described in detail by Soendergaard et al [18]. Briefly, eight kidney pairs were taken from eight brain-dead donor pigs (60-64 kg) and transplanted into sixteen bilateral nephrectomized recipient pigs (14-16 kg), which were randomly given RIC or non-RIC within each pair. Prior to organ reperfusion, RIC was performed as indicated below. Pigs were surveyed, under anesthesia, for 10 h after transplantation, assessing renal function by GFR, renal plasma perfusion and known renal biomarkers. The study was approved by the National Committee on Animal Research Ethics no. 2008-561-1584 (Animal Experiments Inspectorate, Copenhagen, Denmark) and conducted in accordance with the “Principles of Laboratory Animal Care” (NIH publication Vol. 25, No. 28 revised 1996).

### Sample retrieval and ischemia regimen

Kidney retrieval, transplantation and RIC was performed by experienced transplant surgeons as described previously (**Figure 1**) [18]. Briefly, female Danish Landrace pigs were anaesthetized and placed under mechanical ventilation. Ringer’s acetate was infused to maintain fluid balance and arterial pressure. Brain death was induced by increasing intracranial pressure. Kidneys were extracted, perfused and stored in 5°C cold storage solution. Two recipients were transplanted simultaneously after having their native kidneys removed. A RIC regimen was undertaken whereby the abdominal aorta was clamped above the aortic bifurcation but below the renal arteries in four cycles, with each cycle consisting of 5 min of ischemia and 5 min of reperfusion. Surgeons and investigators were unblinded to the treatment during the experiments. Renal biopsies from kidneys in the RIC and non-RIC groups were taken prior to termination after 10 hours reperfusion and snap frozen in liquid nitrogen for transcriptomic and proteomic analysis.

**Figure 1.**
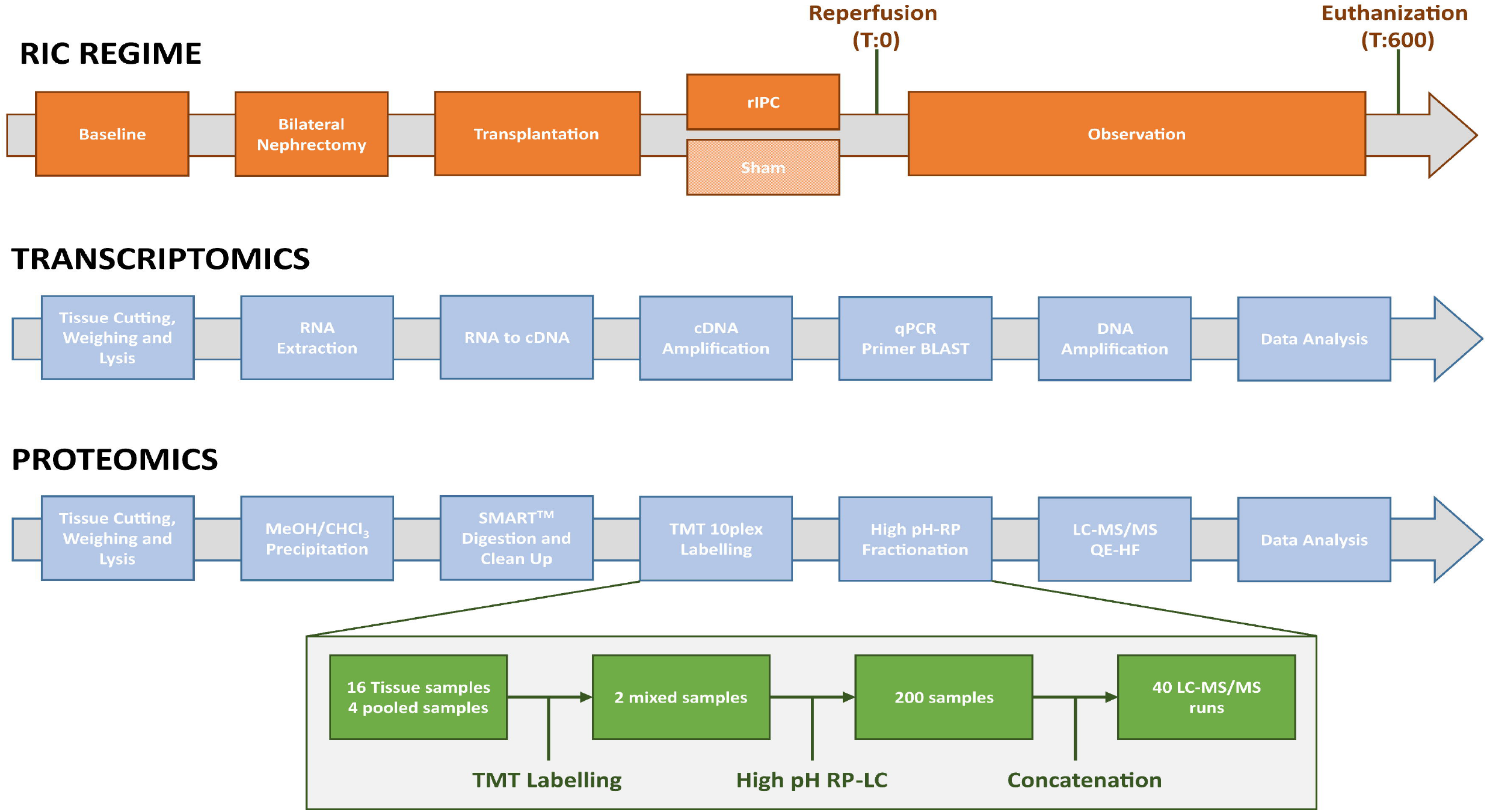
Overview of transcriptomic and proteomic workflows. **A**) For transcriptomics, tissue samples were cut, weight and lysed. RNA was extracted, followed by cDNA library preparation, and the difference in transcript level determined between non-RIC and RIC groups. For selected targets of interest, qPCR was undertaken. **B**) In the proteomics arm, proteins were extracted, reduced, alkylated and subjected to methanol/chloroform extraction, digested with trypsin, cleaned-up and subjected to 10plex TMT labelling. Labelled peptides were separated by high pH-RP, prior to analysis by LC-MS/MS. The green boxes outline our sample pooling and fractionation strategy. A total of 16 tissue samples (and 4 pools) were labelled across two TMT kits, with 8 RIC and non-RIC pairs and 2 pools per 10plex kit. Each TMT experiment was subsequently separated into 100 fractions, which were then concatenated down into 20 fractions. This resulted in a final total of 40 samples for injection into the mass spectrometer.

### Transcriptomics analysis

#### Sample preparation

Sixteen tissue samples from the kidney medulla and cortex (weighing approximately 20 mg) were lysed using a Precellys 24 homogenizer (Bertin Instruments) in 350 μL of RNeasy Mini kit RLT lysis buffer. RNA was then extracted using RNeasy Mini kit (Qiagen, cat number: 74104) as per manufacturer’s instructions. Purified RNA was eluted in 40 μL RNase-free H_2_O and the approximate concentration evaluated using a NanoDrop ND-1000 spectrophotometer. Approximately, 450 ng of purified RNA was converted to cDNA by using high-capacity reverse transcription kit (Applied Biosystems, cat number: 4368814). For each reaction, a mix of 2 μL 10x RT buffer, 0.8 μL 25x dNPT mix, 2 μL 10x RT random primers, 1 μL Multiscribe reverse transcriptase, 3.2 μL nuclease free H_2_O and 11 μL of RNA (40.9 ng/μL) was prepared. This was then primed at 25°C for 10 min, incubated at 37°C for 120 min and inactivated at 85°C for 5 min in a thermal cycler (VWR UNO 96). cDNA was then stored at −20°C until further use. Transcriptomics analysis was performed on ~50⍰fmol of the sample library and sequenced on a NextSeq 500 sequencer (Illumina) achieving, ~30 million reads in total **(Supplementary Table 1)**. An overview of the transcriptomics workflow is given in **Figure 1**.

For transcriptomics data analysis, FASTQ files were converted to Binary-sequence Alignment Format (BAM) files using HISAT2 (v2.1.0) and Samtools (v1.3). The DESeq2 R package (Bioconductor) was used for normalization, visualization, and differential analysis of the data. Furthermore, BAM files were also imported into Perseus software (v1.6.0.2) for comparative analysis, and genome annotation was performed using the *Sus scrofa* FASTA cDNA database (http://www.ensembl.org). Reads per kilo per million (RPKM) values were calculated by a normalization step dividing by the sum (Normalization→Divide), followed by dividing normalized values by gene length, multiplying by 109, and taking the log_2_ values. The transcriptomics dataset was deposited in ArrayExpress (AE) at EBI under the accession number E-MTAB-10872 [21].

#### q-PCR

For q-PCR, primers for selected genes of interest were designed using Primer BLAST (see Table 1 for sequences), parameters included spanning exon-exon junctions, a target size of 200 bp and a maximum product size of 150. A Beta-Actin control gene was used as a reference [22]. A total of 20 μg of cDNA was used for each 20 μL reaction, in combination with 200 nM of each forward and reverse primer (Invitrogen) and Fast SYBR Green Master Mix (Applied Biosystems, cat number: 4385612), according to the manufacturer’s instructions. The reaction plates were run on a Roche Lightcycler 480, with an initial pre-incubation of 37°C for 30 sec, then denaturing at 95°C for 3 sec, followed by annealing at 60°C for 30 sec. Denaturing and annealing was repeated for 40 runs before conducting melt curve analysis. Using Roche Lightcycler analysis software, the noise band was altered to exclude any background noise whilst staying within the lower third of the linear section of the curve to gain initial Ct values. Average Ct values for Beta-Actin were then subtracted from average Ct values of target genes to get dCt. RIC vs non-RIC groups and different time points were then compared using 2^-(x-y) to obtain relative expression levels and plotted on a box-dot graph.

**Table 1:**
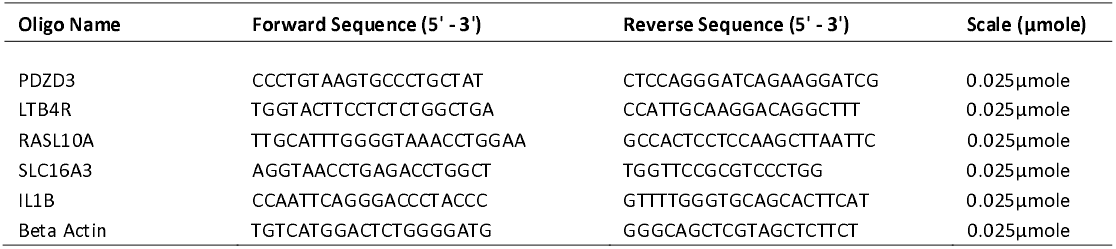
List of gene names and primer sequences used to conduct qPCR. A total of five genes were selected for qPCR validation of transcriptomics results, with beta actin serving as a control.

### Proteomics analysis

#### Sample preparation

In accordance with the transcriptomics study, sixteen tissue samples (weighing approximately 20 mg) were lysed using a Precellys 24 homogenizer (Bertin Instruments) in 30 μL RIPA and 6M urea lysis buffer per 1 mg of tissue. Samples were reduced with 200 mM tris(2-carboxyethyl)phosphine (TCEP) at 55°C for 1 hour and alkylated with 375 mM iodoacetamide (IAA) for 30 min at room temperature. Following this, protein precipitation via methanol/chloroform precipitation was performed. The resulting pellet was dried and resuspended in 50 μL triethylammonium bicarbonate (TEAB) and the protein concentration determined using a BCA assay (Pierce). In total, 100 μg of protein was digested by trypsin using the SMART method (Thermo Scientific). In brief, each sample was loaded into a SMART digestion tube (Thermo Fisher Scientific, Cat no 60109-103) containing 150 μL of SMART digestion buffer. These were then incubated at 70°C at 1400 rpm for 2 hours on a heat shaker (Eppendorf) and spun at 2500 x g for 5 min, with collection of the supernatant. Samples were de-salted using SOLAμ SPE plates (Thermo Scientific, Cat no 60109-103). Columns were equilibrated using 100% acetonitrile (ACN), then 0.1% trifluoroacetic acid (TFA). The samples were then diluted with 1% TFA and pulled through the column using a vacuum pump. They were then washed with 0.01% TFA and eluted in 100 μL 65% acetonitrile. Eluates were dried using a vacuum concentrator (SpeedVac, Thermo Scientific) and resuspended in 100 μL of 100 mM TEAB for Tandem Mass Tag (TMT) labelling.

#### TMT labelling and high pH fractionation

TMT 10plex reaction groups were used to label the digested peptides. Groups comprised of 8 samples (4 sets of RIC and non-RIC) and two pools; one undiluted, and one diluted 1:5, with the pools allowing us to assess the dynamic range of the measurement. For labelling, 41 μL of TMT label was added to each sample and incubated for 1 hour at room temperature. To quench the reaction, 8 μL of 5% hydroxylamine was added to each sample and incubated at room temperature for 15 minutes. Equal volumes of each RIC and non-RIC sample were then pooled into their respective groups to generate a RIC and non-RIC “pooled” sample, and de-salted using Sep-Pak C18 columns. The resulting eluent was dried using a vacuum concentrator (SpeedVac, Thermo Scientific) and resuspended in 120 μL of buffer A (98% MilliQ-H2O, 2% acetonitrile, 0.1% TFA). The two combined TMT 10plex experiments were then pre-fractionated using high pH 10.0 reversed-phase liquid chromatography (HPLC) into a total of 100 fractions. Fractions were subsequently concatenated into a total of 20 samples and submitted for LC-MS/MS analysis. The proteomic sample preparation strategy is summarized in **Figure 1**.

#### Phosphoproteomics

For phosphopeptide enrichment, 50 μg of protein was removed from each of the 16 tissue samples and pooled into their respective RIC and non-RIC groups. Samples were reduced, alkylated, digested with trypsin, and desalted as above. Peptides (~400 μg) were eluted from Sep-Pak cartridges in 500 μL buffer B (65% ACN, 35% MilliQ-H2O, and 0.1% TFA). Phosphopeptide enrichment was performed using an Immobilized Metal Affinity Chromatography (IMAC) column with the aid of a Bravo Automated Liquid Handling Platform (Agilent) according to manufacturer’s instructions.

#### Mass Spectrometry analysis

Liquid chromatography tandem mass spectrometry (LC-MS/MS) analysis was undertaken using a Dionex Ultimate 3000 nano-ultra reversed-phase HPLC system with on-line coupling to a Q Exactive High Field (HF) mass spectrometer (Thermo Scientific). Samples were separated on an EASY-Spray PepMap RSLC C18 column (500 mm⍰×⍰75 μm, 2 μm particle size; Thermo Fisher Scientific) over a 60 min gradient of 2-35% ACN in 5% dimethyl sulfoxide (DMSO), 0.1% FA, and the flow rate was⍰~⍰250 nL/min. The mass spectrometer was operated in data-dependent analysis mode for automated switching between MS and MS/MS acquisition. Full MS survey scans were acquired from 400-2000 *m/z* at a resolution of 60,000 at 200 *m/z* and the top 12 most abundant precursor ions were selected for high collision energy dissociation fragmentation. MS2 fragment ion resolution was set to 15,000.

#### Data analysis

MS raw data files were searched using Proteome Discoverer (v2.3; total proteome) and MaxQuant (v1.5.14; phosphoproteome) software packages. Search parameters included the allowance of two missed trypsin cleavages. Carbamidomethylation (C) was set as a fixed modification, while TMT6plex (N-term, K), oxidation (M), deamidation (N, Q), and phosphorylation (STY) were set as variable modifications where appropriate. The data was searched against porcine protein sequences using the *UPR_Sus scrofa* fasta file (as of August 2019), along with the corresponding decoy reverse database. Only unique and razor peptides were used for quantitation.

#### Statistical analysis

TMT 10plex quantitation and data analysis was performed in Perseus software (v1.6.0.2) which filtered out contaminant and false positive identifications (decoys). The data were log_2_ transformed, and replicates were grouped into RIC and non-RIC. Missing values were imputed from the normal distribution and all samples were normalized by median subtraction. Hierarchical cluster analysis was performed on all the proteins identified by LC-MS/MS and visualized as heat maps using the Pearson correlation algorithm. Volcano plots were generated by applying parametric Student’s t-tests between RIC and non-RIC controls. A Christmas tree plot was generated for the phosphoproteomics data where only one replicate was acquired. The mass spectrometry proteomics data have been deposited to the Proteome×change Consortium via the PRIDE [23] partner repository with the dataset identifier PXD025273. Gene Ontology and biological pathway enrichment analysis was performed using STRING protein-protein interaction networks & functional enrichment analysis (https://string-db.org/).

## RESULTS and DISCUSSION

We first performed RNA sequencing on renal cortex and medulla tissues of kidney grafts to quantify changes at the transcriptome level (**Figure 1**). A total of 19,220 transcripts were successfully quantified across RIC and non-RIC groups (**Supplementary Tables 1 and 2**). In total, 33 transcripts had at least a two-fold change in abundance between groups, at a *p*-value of ≤ 0.05 (**Figure 2, Supplementary Table 3**). The most notable of these included heat shock protein beta-1 (HSPB1), monocarboxylate transporter 4 (SLC16A3), C-X-C motif chemokine (CXCL13), C-C motif chemokine 2 (CCL2), interleukin-1 receptor type 2 (IL1R2), interleukin-1 beta (IL1B), leukotriene B4 receptor 1 (LTB4R), Na(+)/H(+) exchange regulatory cofactor NHE-RF4 (PDZD3), and Ras-like protein family member 10A (RASL10A), all of which were significantly down-regulated following RIC. Moreover, biological pathway analysis revealed a reduction in the transcripts of genes associated with regulation of inflammation following RIC as compared to their non-RIC counterparts (**Figure 2, Supplementary Figure 1**). Transcript changes were confirmed by performing qPCR on a selected panel of targets which were observed to be dysregulated in RIC in either our dataset, or as previously described in literature. We measured five genes in total in a pooled sample of each group, including IL1B, LTB4R, PDZD3, SLC16A3, and RASL10A (**Supplementary Table 4**). We observed a slight increase in RIC induced SLC16A3 transcripts, but overall, none of the genes quantified reached statistically significance between RIC and non-RIC, with all having a *p*-value of > 0.2 (**Figure 3**).

**Figure 2.**
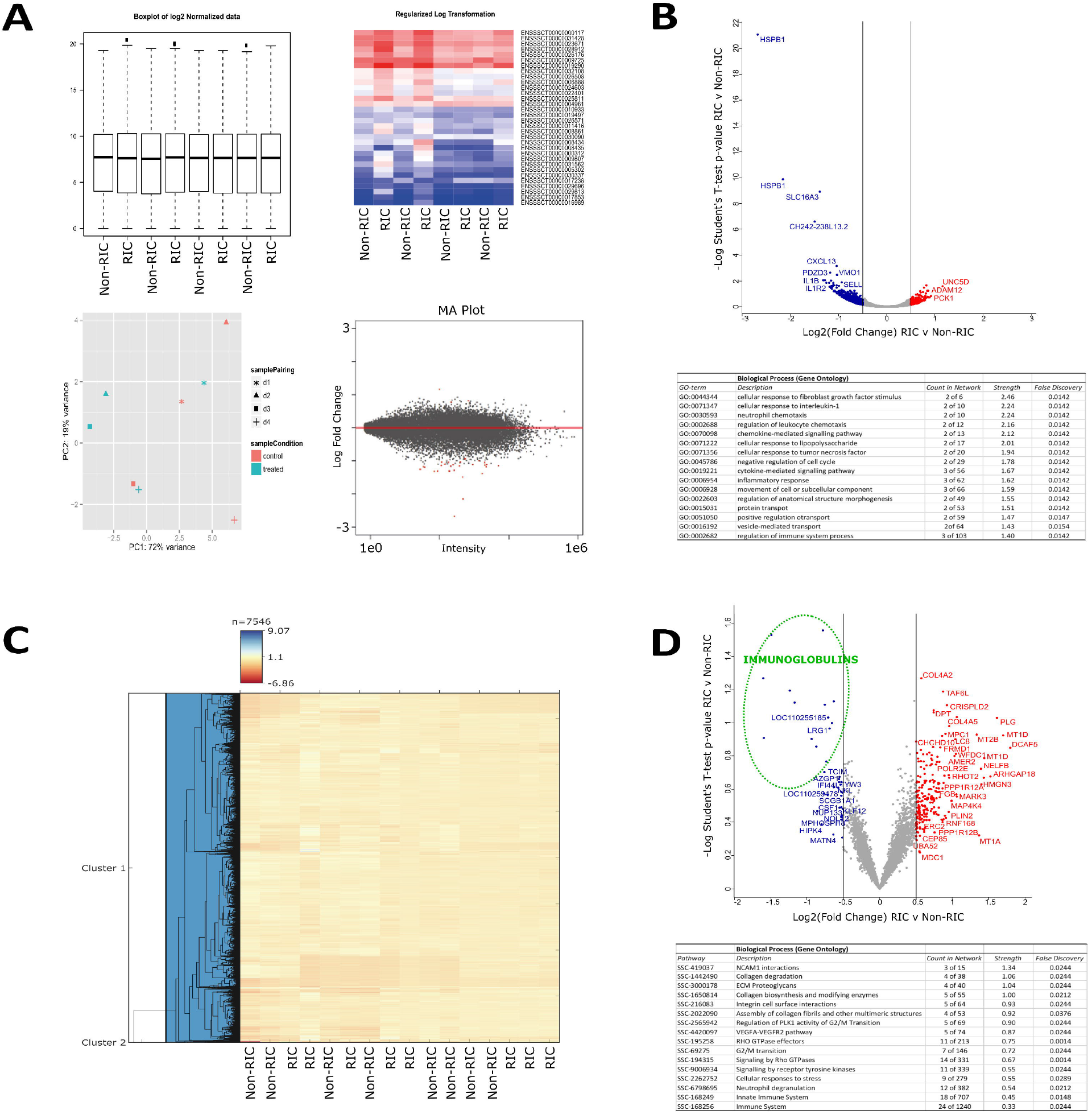
Transcriptome and proteome changes by RIC. **A**) The top left-hand corner displays Box plots of log_2_ normalized data. The heat map displays the regularized log-transformed data for the top differentially expressed transcripts. The PCA plot visualizes the regularized log-transformed data which minimizes differences between samples for rows with small counts, and which normalizes with respect to library size. Finally, the MA plot shows the log_2_ fold changes attributable to a given variable over the mean of normalized counts. Red points have adjusted p-value less than 0.05. **B**) Volcano plot displaying differential transcriptome expression. The x-axis measures expression difference by log_2_(fold change), and the y-axis indicates statistical significance by −log_10_(*p*-value). Down-regulated species are coloured blue, while their up-regulated counterparts are coloured red. Enriched biological processes (Gene Ontology) determined from differentially regulated transcripts (displayed by their gene symbol) are listed underneath the plot. **C)** Hierarchical clustering analysis of RIC and non-RIC proteomes. **D)** Volcano plot displaying differential proteome expression. Down-regulated immunoglobulins of interest are circled in green.

**Figure 3.**
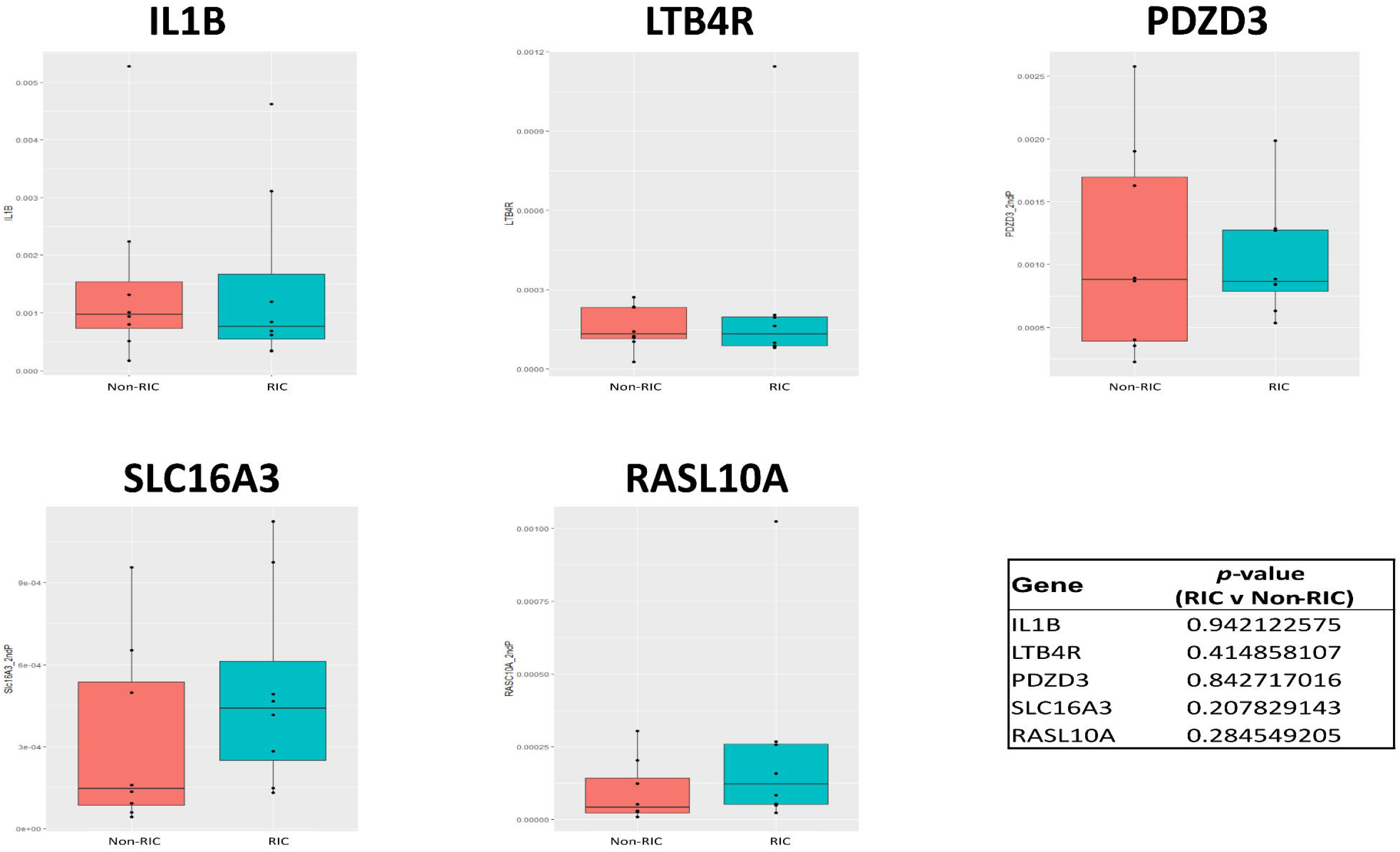
Minimal transcriptional alteration by RIC. No statistically significant difference was observed for any of the selected qPCR targets (IL1B, LTB4R, PDZD3, SLC16A3, and RASL10A) between RIC and non-RIC groups, with all *p*-values > 0.2.

Following the analysis of the kidney donor biopsies at the transcriptome level, we extended the study to measure changes between the kidney tissue proteomes of the respective RIC and non-RIC groups. Our quantitative proteomic strategy (**Figure 1**) successfully quantified a total of 7,546 proteins across all samples (**Supplementary Table 5**). Hierarchical clustering analysis revealed no clear discrimination between RIC and non-RIC groups at the proteome level (**Figure 2A**). Similarly, and in agreement with the transcriptomics results, the majority of proteins were found to be unchanged between groups. To determine if we could identify trends in the data, we took those proteins with the largest (≥ log_2_0.5), albeit non-significant, fold changes forward for further analysis. In this context, 252 proteins were deemed to be differentially expressed between RIC and non-RIC groups (**Supplementary Table 6**). Mirroring the transcriptomics results, a dampening of immune response proteins was observed in RIC versus non-RIC tissue, whereby the largest fold changes were observed in approximately 15 Ig-like domain-containing proteins (**highlighted in green in Figure 2B**). Attenuation of proteins involved in the inflammatory response associated with IRI could be one of the primary beneficial factors that could contribute to RIC renoprotection. This dampening may serve as a natural defense mechanism against the destruction caused by IRI. Despite this, the inflammatory response generally associated with renal transplantation, in the form of apoptotic cell death, macrophage and neutrophil infiltration, and anti-inflammatory cytokine production, was found to be unaffected by RIC in the pigs studied [24]. We performed pathway enrichment analysis on the differentially expressed proteins. Unfortunately, the Ig-like species described above could not be included due to lacking a mature porcine gene entry (**Supplementary Figure 2**). Mirroring the human CONTEXT study [19], there was an accumulation of muscle-derived and extracellular matrix assembly proteins in the renal tissue following RIC, including the collagens COL1A2, COL2A1, COL4A2, COL4A, COL6A3, fibrillin and fibulin (**Figure 2, Supplementary Figure 2**), although no tourniquet on a limb but rather clamping of the aorta was used in the present porcine study. Additionally, upregulation of factors associated with innate immunity such as lysosome-associated membrane glycoprotein 1 (LAMP1), neutrophil cytosol factor 2 (NCF2), leucine-rich alpha-2-glycoprotein (LRG1), myeloperoxidase (MPO), and properdin (CFP) were observed, suggesting the technique provoked the activation of tissue remodeling cascades within the organ itself. Upregulation of these factors provide a favorable environment for wound-healing and repair, and collagen scaffolds have been shown to enhance survival of transplanted cardiomyoblasts and improve function in ischemic rat hearts [25]. Increased collagen production has also been associated with a response to the hypoxic or anoxic conditions inherent to surgery, ultimately leading to adhesion development [26]. It is, however, unclear if these proteins provide a protective mechanism in RIC, or just the consequence of tissue damage which could lead to fibrosis. Furthermore, we identified an up-regulation of several components of the blood coagulation cascade, including Vitamin K-dependent protein C (PROC), Fibrinogen beta chain (FGB), and Plasminogen (PLG). Interestingly, SLC16A3, IL1B, LTB4R, RASL10A, and CXCL13, which were found to have some of the largest changes at the transcript level, were not detected at the proteome level, while HSPB1, LTB4R, and PDZD3 transcripts were differentially abundant, but found to be unchanged between non-RIC and RIC groups at the protein level.

To gain more insight into the signaling dynamics underlying these molecular changes in kidney tissue, we performed phosphoproteomics to compare a global pool of all 8 RIC samples to that of the 8 non-RIC controls. We identified 3,524 phosphosites across both groups, from a total of 3,626 proteins (**Supplementary Table 7**). No substantial differences were identified in the phosphoproteome between RIC and non-RIC groups, indicating little steady-state effects of RIC on kidney tissue at the indicated time point (**Supplementary Figure 3**).

To determine if a joint analysis could reveal useful insights that could not be deciphered from individual analysis alone, we integrated the transcriptomic and proteomic datasets (**Figure 4A**). Once again, the Ig-like proteins that were found to be down-regulated in RIC in the proteomics dataset alone (**Figure 2B**) could not be included in this type of analysis due to a lack of matching cognate sequences at the transcriptome level. A low correlation was observed between the two datasets, with a Pearson correlation coefficient of 0.017. We subsequently performed 2D Annotation Enrichment Analysis in an effort to identify pathways which were simultaneously up- or down-regulated in both datasets [27]. Up-regulated biochemical pathways in RIC versus non-RIC and common to both datasets included muscle tissue development/remodeling, extracellular matrix disassembly, ubiquitin-specific protease activity, and steroid hormone receptor activity (**Figure 4**). Conversely, down-regulated pathways in RIC versus non-RIC and common to both included kidney development, vasodilation, epidermal growth factor receptor signaling and the regulation of myoblast differentiation.

**Figure 4.**
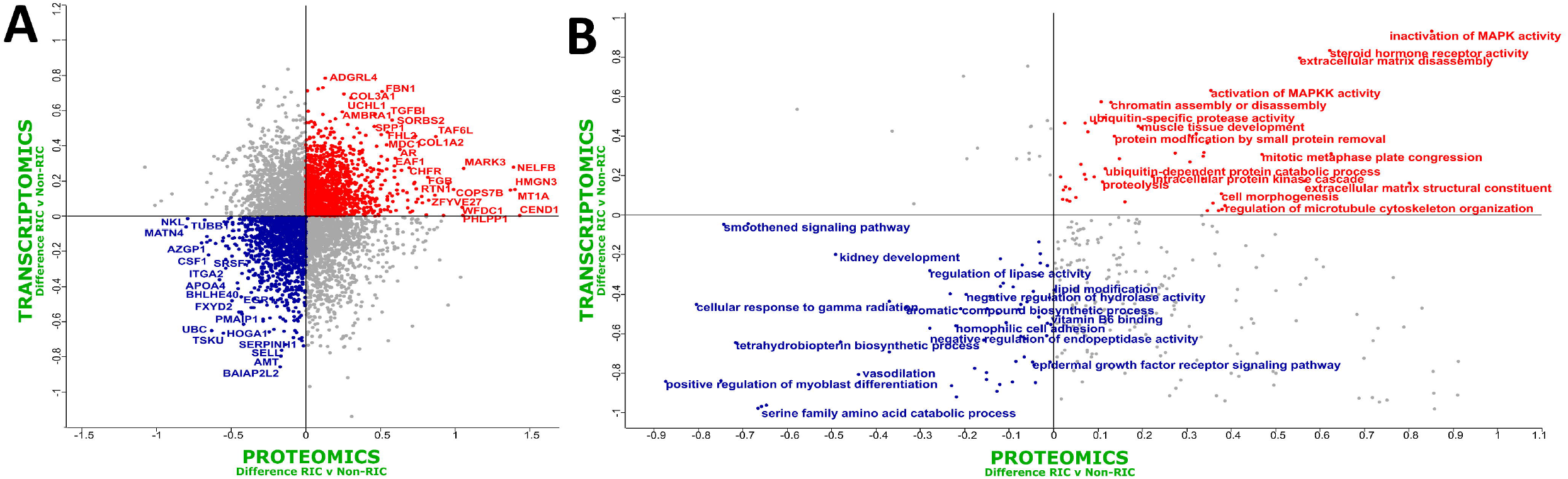
Integrated Omics reveals RIC induced tissue leakage and reduced inflammation. A) Scatter plot of the *Log_2_-fold-change* between RIC and non-RIC groups of the proteomics on the x-axis, versus the transcriptomics on the y-axis. Species which are down-regulated in both datasets are coloured blue, while their up-regulated counterparts are coloured red. A low Pearson correlation coefficient of 0.017 was observed. B) 2D Annotation Enrichment Analysis generated using transcripts and proteins which were commonly dysregulated across the two datasets.

A possible explanation for the absence of substantial and significant proteomic and transcriptomic changes may be attributed to the condition of kidney tissue used in this study. The tissue had been exposed to multiple insults, including brain death, periods of cold and warm ischemia and reperfusion. This makes it increasingly difficult to fully dissect the (perhaps comparatively minor) effects of RIC, despite a more aggressive RIC regimen than that used in the human setting. A limiting factor of this study concerning -Omics profiling is the number of available samples, and the timepoint at which the analyzed biopsies were taken after a 10-hour observation phase. This may be too early of a time point to measure a significant change in protein expression, which is possibly reflected in the lack of change identified at the proteomic depth achieved in this study, although the transcriptome profile would not support this notion. Ideally, there would be an observation phase of several days with multiple biopsies. Finally, with the disappointing clinical effects of RIC, the positive effects of RIC regarding GFR and graft reperfusion in the related pig study could reflect a type 1 error or generally reflect a subtle phenomenon observed in this animal model [18].

## CONCLUSIONS

In summary, RIC of the pig recipient before kidney graft transplantation with reperfusion, similar to what was observed in human counterparts, did not result in substantial molecular changes/perturbations in the graft compared to non-RIC controls. The data did suggest, however, that the administered RIC caused a potentially detrimental accumulation of muscle proteins in the kidney graft tissue, and also triggered a subtle interplay between the innate and adaptive immune response, with a dampening of the latter. Ultimately, however, our discovery studies showed no major differences in the transcriptomes and proteomes of porcine kidney transplant tissue following RIC, despite excellent depth of coverage in each. Our results, therefore, support the conclusion that the RIC protocol used does not substantially protect against ischemia/reperfusion injury.

## LIST OF ABBREVIATIONS

AKI: Acute Kidney Injury
ATP: adenosine triphosphate
GFR: Glomerular Filtration Rate
IRI: Ischemic/Reperfusion Injury
RIC: Remote Ischemic Preconditioning
TMT: Tandem Mass Tag

## DECLARATIONS

### ETHICS APPROVAL AND CONSENT TO PARTICIPATE

Kidney graft tissue samples were taken from the porcine RIC study [18].

### CONSENT FOR PUBLICATION

All authors have read the manuscript and given their consent for publication.

### AVAILABILITY OF DATA AND MATERIALS

The mass spectrometry raw data files have been deposited under PRIDE Submission PXD025273. The transcriptomics dataset was deposited in ArrayExpress (AE) at EBI under the accession number E-MTAB-10872.

### AUTHORS CONTRIBUTIONS

DOB, AT, HH, EP, XY, PS, KR and NS performed the experiments. BJ, RJP and BMK conceptualized the study. DOB, AT, BJ, RJP and BMK wrote the manuscript. DOB, AT, HH, EP, PS, KR and NS analyzed the data. All authors read and approved the final manuscript.

## ACKNOWLEDGEMENTS

The authors would like to thank members of the Kessler, Jespersen, and Ploeg research groups for helpful discussions and technical support.

## FIGURE LEGENDS

**Supplementary Figure S1. *Transcriptomic pathway enrichment analysis reveals subtle alteration of tissue inflammation by RIC*.** All transcripts (n=33) which were found to be dysregulated in RIC versus non-RIC controls were searched against the *Sus scrofa* database in STRING. Only interactions of the highest confidence (scores > 0.90) were included in the analysis. Genes associated with immune regulation are highlighted, with those linked to interleukin biology coloured red, while cytokines are coloured purple.

**Supplementary Figure S2. *Proteomic pathway enrichment analysis uncovers RIC induced tissue leakage and altered inflammation*.** Proteins (n=252) which were found to have the greatest dysregulation in RIC versus non-RIC controls were searched against the *Sus scrofa* database in STRING. Only interactions of the highest confidence (scores > 0.90) were included in the analysis. Proteins/genes of interest are highlighted, with proteins associated with muscle and ECM coloured in blue, blood coagulation in green, and factors associated with innate immunity coloured red.

**Supplementary Figure S3. *Kidney tissue phosphoproteomics unaffected by RIC*.** A) Abundance plots indicate no significant difference in the phosphoproteome between RIC and control groups. B) A Christmas tree plot, with Significance B thresholds colour coded. Minor changes were observed between groups.

**Supplementary Table 1: *Transcriptome sequence alignments*.** An average sequence alignment of 82% was achieved across all samples

**Supplementary Table 2: *List of transcripts identified by transcriptomics*.** A total of 19,220 transcripts were successfully identified across groups.

**Supplementary Table 3: *Transcripts differentially expressed between RIC and Non-RIC groups*.**

**Supplementary Table 4: *qPCR validation on a panel of targets from discovery transcriptomics*.** IL1B, LTB4R, PDZD3, SLC16A3, and RASL10A were quantified by qPCR. We observed a slight increase in RIC induced SLC16A3 transcripts, but overall, none of the genes quantified reached statistically significance between RIC and non-RIC, with all having a *p*-value of > 0.2

**Supplementary Table 5: *List of proteins identified by proteomics*.** A total of 7,546 proteins were successfully identified across groups.

**Supplementary Table 6: *Proteins differentially expressed between RIC and Non-RIC groups*.**

**Supplementary Table 7: *List of proteins and phosphosites identified by phosphoproteomics*.** A total of 3,524 phosphosites were successfully identified across groups.

