## Supplementary Figures for "Integrative Omics Reveals Subtle Molecular Perturbations Following Ischemic Conditioning in a Porcine Kidney Transplant Model"

### Slide 1
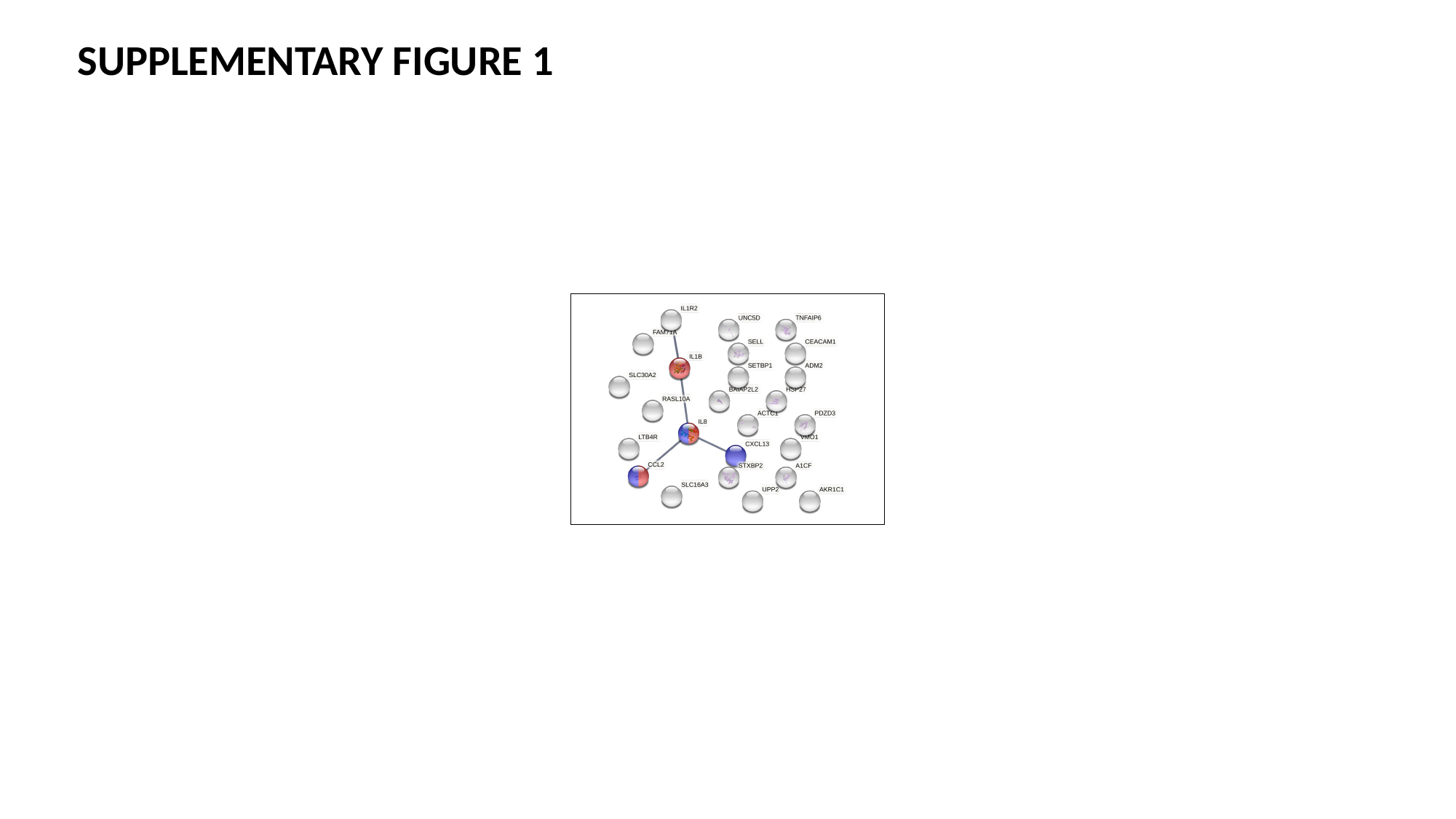

SUPPLEMENTARY FIGURE 1

### Slide 2
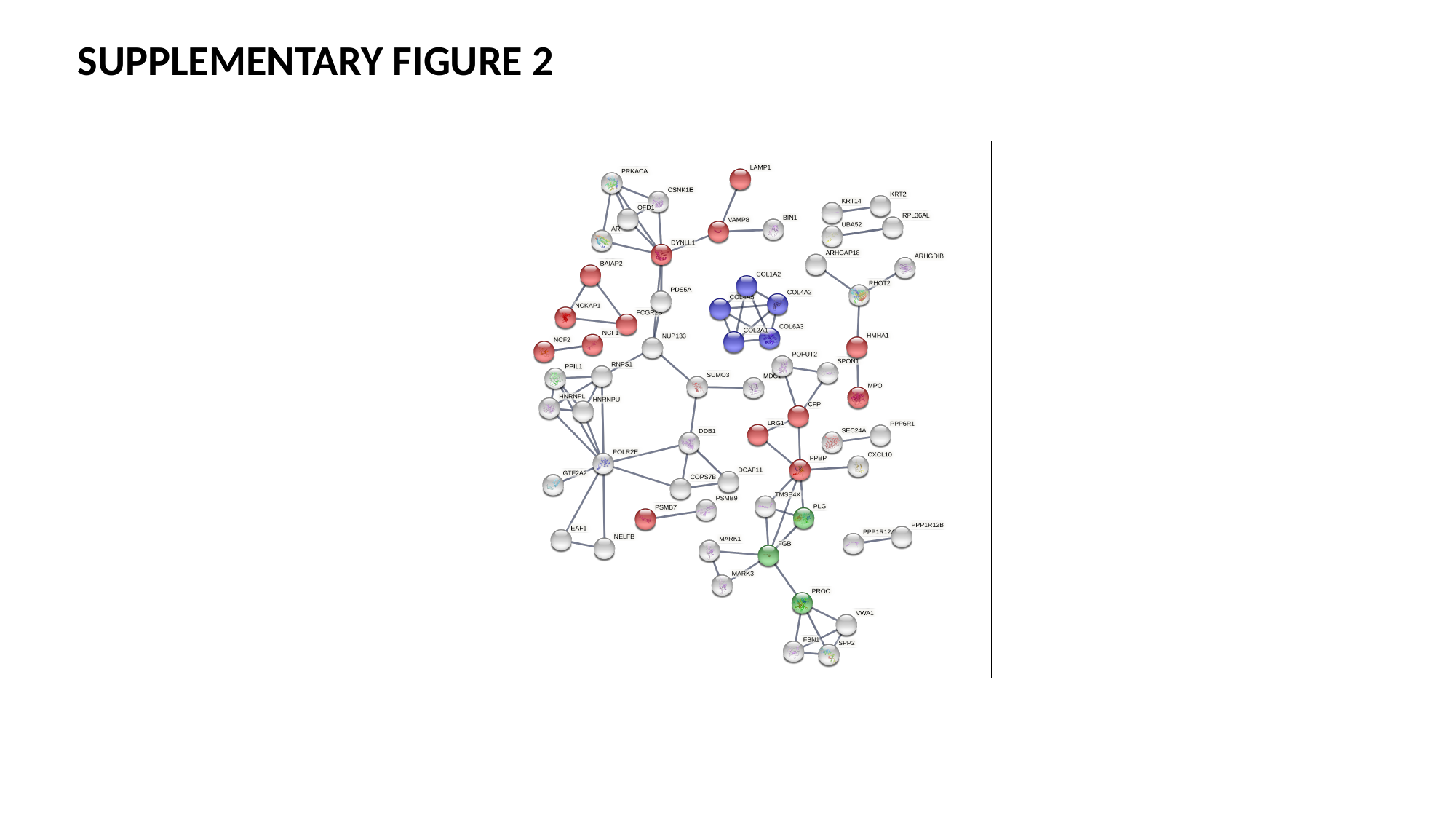

SUPPLEMENTARY FIGURE 2

### Slide 3
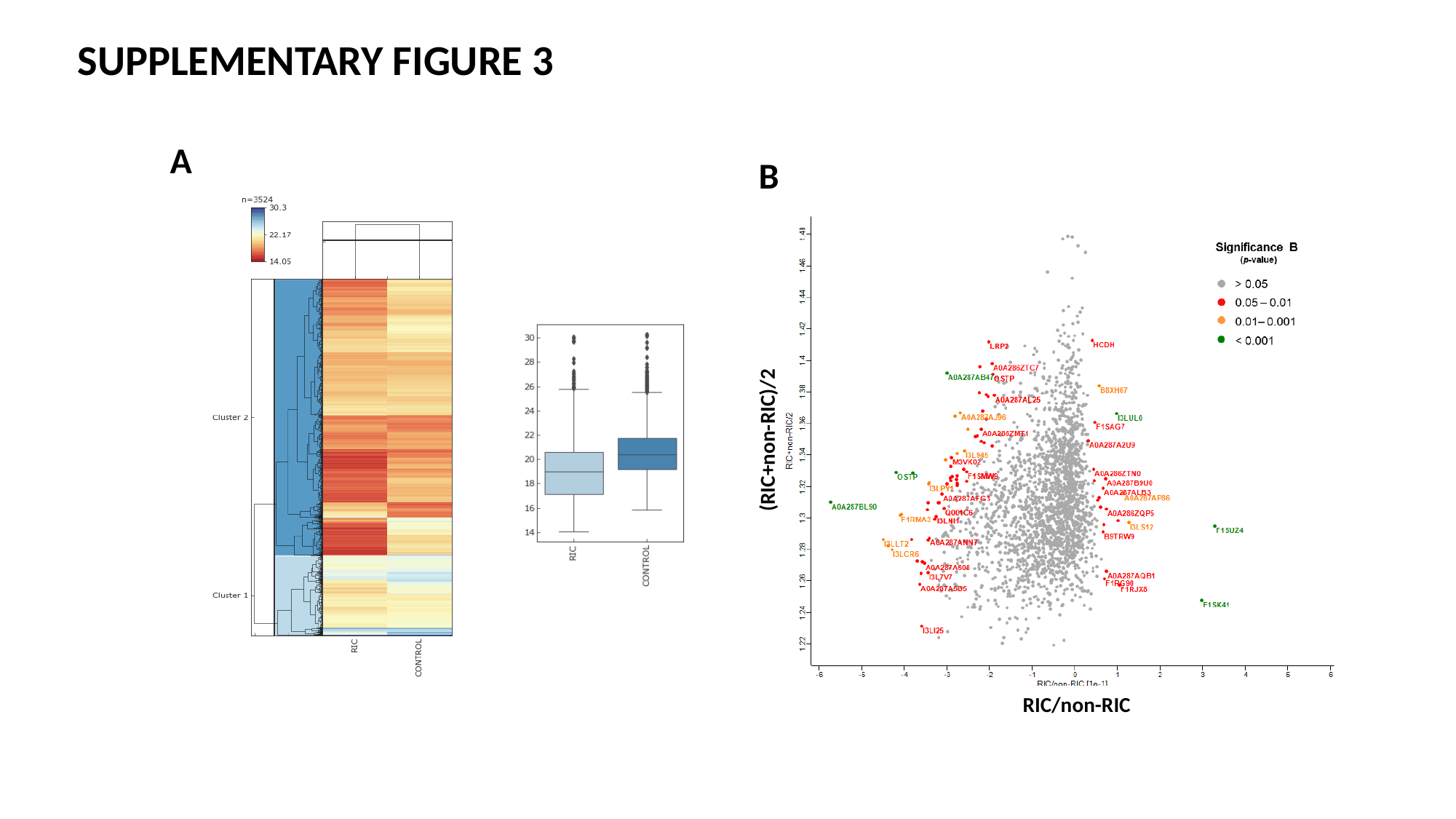

SUPPLEMENTARY FIGURE 3
A
B
(RIC+non-RIC)/2
RIC/non-RIC
